# A multi-scenario risk assessment strategy applied to mixtures of chemicals of emerging concern in the River Aconcagua basin in Central Chile

**DOI:** 10.1101/2023.08.23.554257

**Authors:** Pedro A. Inostroza, Sebastian Elgueta, Martin Krauss, Werner Brack, Thomas Backhaus

**Author notes:** Corresponding author: Pedro A. Inostroza.

## Abstract

Streams and rivers are characterised by the presence of various chemicals of emerging concern (CECs), including pesticides, pharmaceuticals, personal care products, and industrial chemicals. While these chemicals are found usually only in low (ng/L) concentrations, they might still harm aquatic life and disrupt the ecological balance of aquatic ecosystems due to their high ecotoxicological potency. Environmental risk assessments that account for the complexity of exposures are needed in order to evaluate the toxic pressure of these chemicals, which also provide suggestions for risk mitigation and management, if necessary. Currently, most studies on the co-occurrence and environmental impacts of CECs are conducted in countries of the Global North, leaving massive knowledge gaps in countries of the Global South.

In this study, we implement a multi-scenario risk assessment strategy to improve the assessment of both the exposure and hazard components in the chemical risk assessment process. Our strategy incorporates a systematic consideration and weighting of CECs that were not detected, as well as an evaluation of the uncertainties associated with Quantitative Structure-Activity Relationships (QSARs) predictions for chronic ecotoxicity. Furthermore, we present a novel approach to identifying mixture risk drivers. To expand our knowledge beyond well-studied aquatic ecosystems, we applied this multi-scenario strategy to the River Aconcagua basin of Central Chile. The analysis revealed that the concentrations of CECs exceeded acceptable risk thresholds for selected organism groups and the most vulnerable taxonomic groups. Streams flowing through agricultural areas and sites near the river mouth exhibited the highest risks. Notably, the eight risk drivers among the 153 co-occurring chemicals accounted for 66-92% of the observed risks in the river basin. Six of them are pesticides and pharmaceuticals, chemical classes known for their high biological activity in specific target organisms.

**Graphical abstract:** 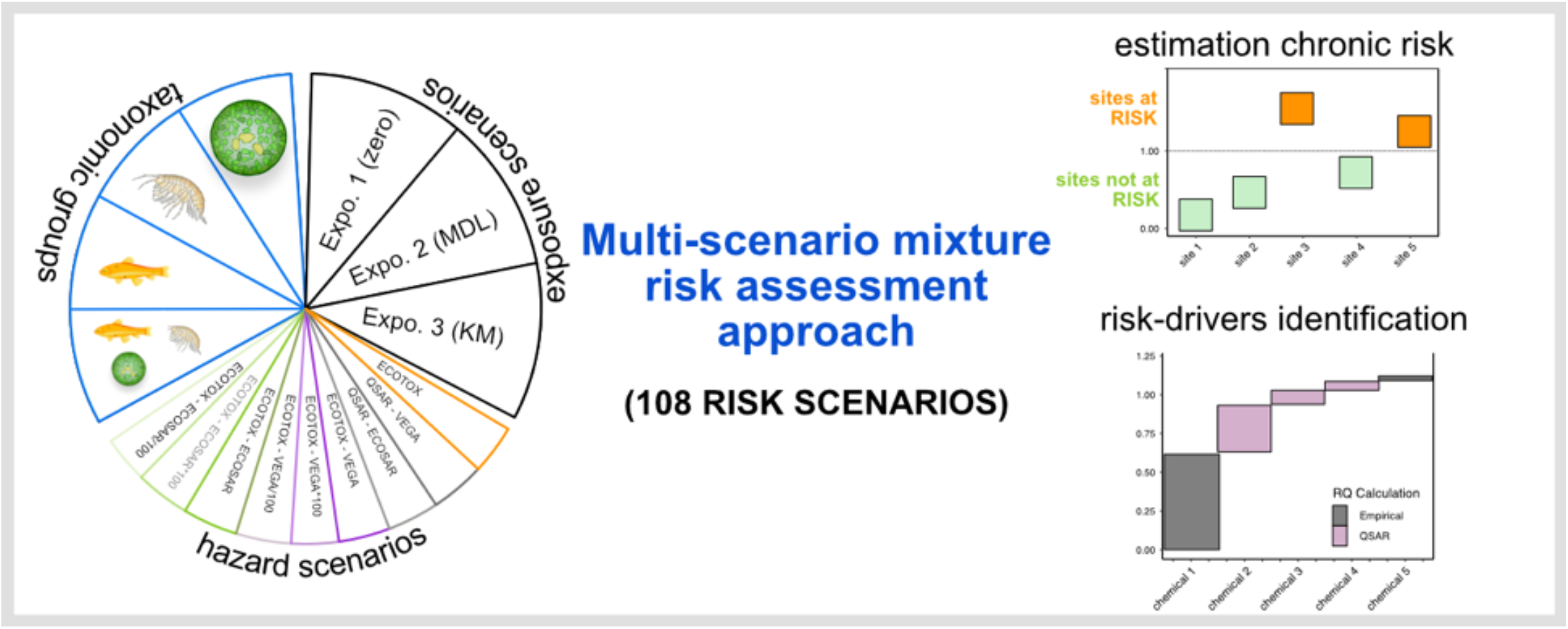

**Highlights:**

- 153 chemicals of emerging concern detected in complex multi-component mixtures.
- 108 possible mixture risk assessment scenarios were investigated.
- Non-detects, QSARs, and experimental ecotoxicological data were integrated for risk assessment.
- 8 chemicals of emerging concern were responsible for driving chronic environmental risks.

## 1. Introduction

Anthropogenic chemical pollution has profound impacts on the ecological status of surface waters at a continental scale (Malaj et al., 2014), and is therefore increasingly recognised as a driving force behind biodiversity loss (Balvanera et al., 2019; Groh et al., 2022; Sigmund et al., 2023). However, especially diffuse pollution is characterised by the presence of complex multi-component mixtures (Finckh et al., 2022; Kandie et al., 2020; Marshall and McCluney, 2021). These mixtures comprise a diverse array of organic chemicals, including pharmaceuticals and personal care products (PPCPs), pesticides, surfactants, industrial chemicals, and transformation products, collectively referred to as chemicals of emerging concern (CECs) (Ankley et al., 2008). CECs are recognised for their harmful effects on aquatic life, even at low environmental concentrations, affecting various organisms ranging from microbes (Drury et al., 2013) to higher vertebrates (Jobling et al., 1998), and exerting influences on genes and the genetic landscape of exposed organisms (Inostroza et al., 2018).

CECs enter streams and rivers through various pathways, including direct discharges from wastewater treatment plants (WWTPs) (Hug et al., 2014), emissions from industrial facilities (Kaewlaoyoong et al., 2018), unintentional runoff from agricultural areas, and accidental spills (Reiber et al., 2021). Despite the low degradability of many CECs, their continuous release into the aquatic environment results in a phenomenon known as “pseudo-persistence” (Boxall et al., 2004; Kolpin et al., 2002), and ultimately in chronic exposures and ecological effects. Conventional WWTPs exhibit limited effectiveness in removing CECs (Eggen et al., 2014), and even advanced technologies, such as phosphorus elimination, nitrification, and denitrification, are ineffective in CEC removal (Neale et al., 2017). Despite numerous studies focusing on CEC concentrations in surface waters, particularly regarding pharmaceuticals (Wilkinson et al., 2022) and pesticides (Chow et al., 2020), significant knowledge gaps persist regarding the co-occurrence and environmental risks associated with CECs, especially in developing countries. These countries often experience the highest CEC concentrations due to inadequate WWTP technologies (Wilkinson et al., 2022) and/or outdated environmental protection frameworks.

Even if all chemicals are present at concentrations below their individual “safe” levels, there may still be an unacceptable risk posed by the mixture (Rudén et al., 2019). The environmental risk associated with these mixtures can be modelled using the concentration addition model (CA), which is widely recommended as an initial precautionary approach for any mixture assessment (Backhaus and Faust, 2012; Kortenkamp et al., 2009; Rudén et al., 2019). CA can be applied to chemical monitoring data and allows exploring different exposure and/or hazard scenarios, in which the reliability and validity of the risk estimates can be systematically explored, for instance, the role of non-detects and Quantitative Structure-Activity Relationships (QSARs), see below. This also permits the identification of the risk-driving chemicals from the considerable number of chemicals that are often found to co-occur.

Chemicals that are included in the monitoring suite, but that are not detected occur at a concentration somewhere between zero and the chemical-analytical limit of detection. The incorporation of non-detects into the risk assessment lacks systematic integration, and only a limited number of studies have examined their potential contribution to the mixture risk (Gustavsson et al., 2017a, 2017b; Rodríguez-Gil et al., 2018). Most studies either simply disregard non-detects altogether or employ substitution methods, which introduce biases in the exposure estimates (Leith et al., 2010).

The CA model requires the ecotoxicological characterisation of every mixture component. However, such data are often lacking for CECs. Those gaps are therefore often bridged by *in silico* methods such as QSARs. However, the resulting uncertainties are often not comprehensively evaluated and integrated into the risk assessment. It is important to note that QSARs for mixtures of chemicals with distinct modes of action are inherently less robust compared to predictions for a single mechanism of action (Escher and Hermens, 2002). Therefore, describing these uncertainties will identify important data gaps and their impact on the final risk estimate.

In recent decades, Chile witnessed a notable increase in agricultural production, resulting in a corresponding increase in pesticide usage (Coria and Elgueta, 2022). The Central Valley, where the River Aconcagua is located, is characterised by agricultural activities, and several studies have investigated the presence of pesticides and their transformation products in surface waters within this region (Climent et al., 2019; Giordano et al., 2011; Montory et al., 2017) (Inostroza et al., accepted - Data in Brief Journal). Although wastewater treatment facilities are widely distributed throughout the country, with a high coverage rate of 96.6% among the urban population (OECD/ECLAC, 2016), it is noteworthy that only two-thirds of urban households are connected to advanced wastewater treatment plants (secondary or tertiary treatment). Additionally, wastewater treatment coverage remains limited in rural areas (OECD/ECLAC, 2016).

Monitoring studies that focus on assessing the environmental risks of organic chemicals in Chile’s aquatic environment, particularly in streams and rivers, are scarce. In comparison, coastal areas have received slightly more attention, with some studies quantifying the presence of antibiotics (Buschmann et al., 2012), endocrine-disrupting chemicals (Bertin et al., 2011), and industrial chemicals (Salamanca et al., 2019). This may reflect the broader situation in countries of the Global South, including Chile, where outdated monitoring programs and inadequate water management frameworks persist (OECD/ECLAC, 2016).

This study, therefore, implements a multi-scenario mixture risk assessment for the River Aconcagua basin, located in Central Chile. It incorporates various exposure scenarios to account for non-detects and CECs with missing empirical ecotoxicological data. We propose a novel strategy for identifying and prioritising mixture risk drivers within complex environmental mixtures.

## 2. Material and methods

### 2.1. Case study area - River Aconcagua Basin

The River Aconcagua (143 km long) is located in Central Chile and its basin drains an area of 7,338 km^2^. The basin is characterised by a Mediterranean climate with warm, dry summers (October to March) and wet, cool winters (May to August) marked by intense and irregular rain (Amigo and Ramírez, 1998). There are half a million residents in this river basin, and it supports 12% of Chile’s national agriculture and 4% of its copper production, respectively (COCHILCO, 2020). Roughly 7.6% of the total river basin is devoted to agriculture, with more than 90% of the cropland (avocado and grapes) concentrated in the Lower and Putaendo sub-basins (Webb et al., 2021). Moreover, ten middle-sized WWTPs, featuring aeration ponds and activated sludge technologies, are located across the basin, serving about 405,000 residents. However, only five of them discharge directly into the main course of the River Aconcagua and the others discharge into its tributaries (Inostroza et al., accepted - Data in Brief Journal).

Surface water samples were collected from nine sampling sites in October 2018 (dry season). Sampling sites were selected based on land use types (e.g., streams and/or rivers running through natural parks, agricultural areas, urban, and mixed land uses). Environmental measured concentrations of CECs in the River Aconcagua Basin along with detailed analytical methodologies are accessible through the zenodo repository (Inostroza et al., 2023) and Inostroza et al., (accepted - Data in Brief Journal), respectively. Additionally, sampling sites are reported as supplementary material in Table S1.

### 2.2. Retrieval and curation of empirical ecotoxicological data

Chronic experimental data were obtained from the US EPA ECOTOXicology Knowledgebase (ECOTOX) version “ecotox_ascii_09_15_2022” (Olker et al., 2022) for all the targeted chemicals. Only data for freshwater organisms and only for chronic exposures were retained. The data were furthermore curated by excluding records that lacked values for exposure durations, measurement endpoints, appropriate units, or a reference for the data source, as well as limit values, were excluded. To ensure uniformity, effect concentrations were normalised to µmol/L. Chronic effect data were identified in accordance with Australian and New Zealand guidelines (Warne et al., 2018), with exposure durations of at least 1, 14, and 21 days for algae, macroinvertebrates, and fish, respectively. All remaining data were recalculated to chronic EC10-equivalents, using the extrapolation factors from (Warne et al., 2018). LC/IC/EC50 values, LOECs and MATC values were divided by 5, 2.5, and 2, respectively. An in-house dataset was used to assign a taxonomic group (i.e., algae, macroinvertebrates, and fish) to each species group based on the reported phyla in ECOTOX and all other data were discarded. For each taxonomic group, the geometric mean was calculated for each chemical. The geometric mean was chosen over the arithmetic mean as it is considered more resistant to the impact of outliers and more suitable for skewed datasets (Leith et al., 2010).

### 2.3. Quantitative Structure Activity Relationships (QSARs)

Limiting the assessment to those chemicals for which chronic experimental data are available results in an underestimation of the mixture risk. Various academic researchers as well as regulatory authorities such as the European Chemicals Agency (ECHA) and the US EPA encourage the use of quantitative structure-activity relationships (QSARs) in order to estimate ecotoxicological properties *in silico*. QSARs were employed to predict the chronic toxicity for algae, macroinvertebrates and fish, for all chemicals detected at least once. Two QSAR platforms, the VEGA HUB (version 1.1.5 48, (Benfenati et al., 2013)) and the Ecological Structure Activity Relationships (ECOSAR) Class Program (version 2.2) were utilised for this purpose. Chemicals were identified via their CAS numbers, and the corresponding SMILES (Simplified molecular-input line-entry system) were then used as a chemical identifier for the QSAR calculations. If multiple predictions were provided by the software, its geometric mean was used for the mixture risk assessment. The predicted toxicities were transformed to µmol/L.

### 2.4. Mixture risk assessment

A mixture risk assessment can be either performed separately for each of the three taxonomic groups (algae, macroinvertebrates, fish) or by accounting for the most sensitive taxonomic group (MST) for each chemical. The MST approach is conceptually similar to first calculating a Predicted No Effect Concentration (PNEC, European Chemicals Agency, 2016) or Environmental Quality Standard (QSfw,eco European Commission, 2018) without applying any assessment factor and then applying Concentration Addition to these values (Gustavsson et al., 2017a).

The environmental mixture assessment was conducted using CA, for details see (Gustavsson et al., 2017a; Rudén et al., 2019; Spilsbury et al., 2020). The mixture risk for a particular taxonomic group (algae, macroinvertebrates, fish), expressed as its risk quotient (RQ_STU_), is defined as follows:

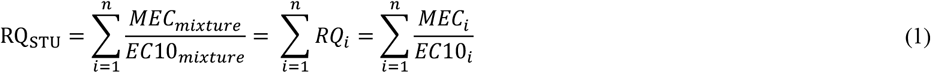

where *MEC*_i_ is the measured environmental concentration of chemical *i* and EC10_i_ denotes the corresponding geometric mean of chronic effect concentrations (EC10-equivalent) of chemical *i* for a particular taxonomic group (algae, macroinvertebrates or fish). The ratio *MEC*_i_/EC10_i_ provides a dimensionless measure of the toxicity contribution of chemical *i*. This approach estimates the mixture risk quotient separately for each taxonomic group.

The ecological risk posed by the mixture of CECs was evaluated on a more integrating ecological level through the application of the concept of the most sensitive taxonomic group (MST) (Backhaus and Faust, 2012; Gustavsson et al., 2017a), in which the mixture risk quotient is defined as:

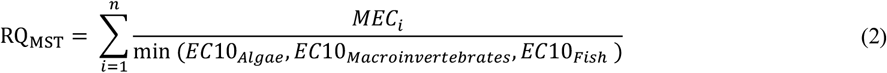

This method corresponds to the summation of fractions of Predicted No Effect Concentrations (PNECs) without using any assessment factors (Backhaus and Faust, 2012; Gustavsson et al., 2017a). In line with the strategy outlined for the environmental risk assessment of industrial chemicals under REACH, we applied the MST approach by combining the data from three taxonomic groups (European Commission, 2011).

For the assessment of risks for each individual taxonomic group as well as the MST, we employed a final assessment factor of 10, again in line with the European guidelines for industrial chemicals, in order to account for the extrapolation from the laboratory to the field situation and to account for the lack of biodiversity considerations in the assessment (European Commission, 2011).

For all these calculations, ecotoxicity data for all detected chemicals are required. Empirical data gaps were bridged by QSARs (see above. However, as QSAR-estimates had a comparatively low accuracy (see results), we included specific scenarios in which we assumed that the QSAR estimates were off by two orders of magnitude (Table 1, see results for a justification on why two orders of magnitude were used as the likely margin of error). All in all, nine different hazard scenarios were included in the assessment (Table 1).

**Table 1.** Hazard scenarios used for environmental mixture assessment.

| Scenarios | Definition |
| --- | --- |
| ToxA | Only experimental chronic toxicity data from US EPA ECOTOX Database |
| ToxB | Only QSAR VEGA HUB chronic predictions |
| ToxC | Only QSAR ECOSAR chronic predictions |
| ToxD | Experimental toxicity data amended with QSAR VEGA |
| ToxE | Experimental toxicity data amended with QSAR VEGA *100 |
| ToxF | Experimental toxicity data amended with QSAR VEGA /100 |
| ToxG | Experimental toxicity data amended with QSAR ECOSAR |
| ToxH | Experimental toxicity data amended with QSAR ECOSAR *100 |
| ToxI | Experimental toxicity data amended with QSAR ECOSAR /100 |

Three exposure scenarios were defined, depending on how non-detects were accounted for:

i. Exposure-Scenario 1: non-detects were set to zero, representing the scenario with the lowest risk that is still compatible with the analytical data.
ii. Exposure-Scenario 2: non-detects were set to their method detection limits (MDLs), representing the scenario with the highest risk that is still compatible with the analytical data.
iii. Exposure-Scenario 3: missing concentration values were estimated using Kaplan-Meier modelling (Gustavsson et al., 2017a; Helsel, 2010), providing the most accurate basis for the risk assessment but not allowing to identify individual risk drivers.

We identified two categories of mixture risk drivers: absolute and relative risk drivers. An absolute risk driver is defined as a compound that contributes to the mixture risk with an RQ of at least 0.02 (i.e., 20% of the acceptable mixture RQ of 0.1), at least at one site. A relative risk driver is a compound which contributes 20% or more of the final RQ-sum, at least at one site (Table 2). Given the considerable uncertainty introduced by bridging data gaps with QSAR estimates (see above), we termed compounds that are not identified as risk drivers but could become one if the QSAR value underestimates the compound’s toxicity by at least 2 orders of magnitude as *potential* mixture risk drivers (Table 2). Actual risk drivers are compounds that should be prioritised for risk mitigation, while *potential* risk drivers are compounds flagged for ecotoxicological testing.

**Table 2.** Mixture risk driver definitions.

|  | Definition used in this study |
| --- | --- |
| Absolute risk driver | A compound with an RQ of at least 0.02. |
| Relative risk driver | A compound that contributes at least 20% to RQ-sum |
| *Potential* absolute risk driver | A compound that is not an absolute risk driver but becomes one if their QSAR-estimated RQ-contribution is increased by a factor of 100. |
| *Potential* relative risk driver | A compound that is not a relative risk driver but becomes one if their QSAR-estimated RQ-prediction is increased by a factor of 100. |

### 2.5. Data analysis

The statistical analyses and data visualization were performed using R version 4.2.2 (R Core Team, 2021). To assess the normality assumption, the Shapiro-Wilk Normality Test was utilised, and if the data deviated from a normal distribution, non-parametric testing was employed. The comparison of quantified environmental concentrations across chemical classes and sampling sites was conducted using the Kruskal-Wallis test (KW) and Dunn’s test, which was implemented in the R-package {dunn.test}(Dinno, 2017), respectively. The Kaplan-Meier adjustment was incorporated in the analysis using the R-package {NADA} (Helsel, 2005). All data curation and data analysis scripts are available on Github (https://github.com/ThomasBackhausLab/Mixture_assessment_analysis).

## 3. Results

### 3.1. Occurrence of CEC mixtures in the River Aconcagua Basin

Detailed tables with detected and quantified CECs, concentrations, and their respective chemical identifiers are published in a separate data paper (Inostroza et al., accepted - Data in Brief Journal) and are available via zenodo (Inostroza et al., 2023). The data reveal the widespread occurrence of CECs in the surface waters of the River Aconcagua basin. From the 861 organic chemicals included in the analysis, 153 chemicals, including PPCPs, pesticides, and industrial chemicals were detected and quantified at least at one site. The industrial chemicals triacetonamine (intermediate and potential degradation product of plastic additives (UV stabilizers)) and benzyl dimethyl ketal (UV photosensitizer) as well as the disinfectant didecyldimethylammonium (DDA) were detected at all sites (Figure S1). The number of detected and quantified chemicals varied across sampling sites. We detected and quantified between 46 and 80 chemicals in tributary streams, between 39 and 71 in the main river course, and only between 18 and 28 at the reference sampling sites. The high number of CECs in tributary streams is likely due to intensive agriculture and the influence of WWTP discharges near the sampling sites. The low number of CECs found at RS1, RS2, and RS3 sites is a result of the lower urbanisation in the region and supports the use of these sites as “reference sites”.

The highest measured environmental concentrations were recorded in the main river course (35,625 ng/L) followed by tributaries (6,038 ng/L), and reference sites (655 ng/L). The highest CEC concentrations (top 20%) are plotted in Figure 1. The artificial sweetener sucralose reached the highest concentrations in the main river course (538-35,625 ng/L) and tributary streams (1,688-6,038 ng/L), most likely as a consequence of its discharge from the WWTPs spread along the river basin. Benzothiazole, a vulcanisation accelerator but also used as a UV stabiliser and pesticide, was found in almost similar concentrations in the reference sites (501-655 ng/L) and tributaries (452-496 ng/L), while the main river course was slightly less exposed (37-345 ng/L).

**Figure 1.**
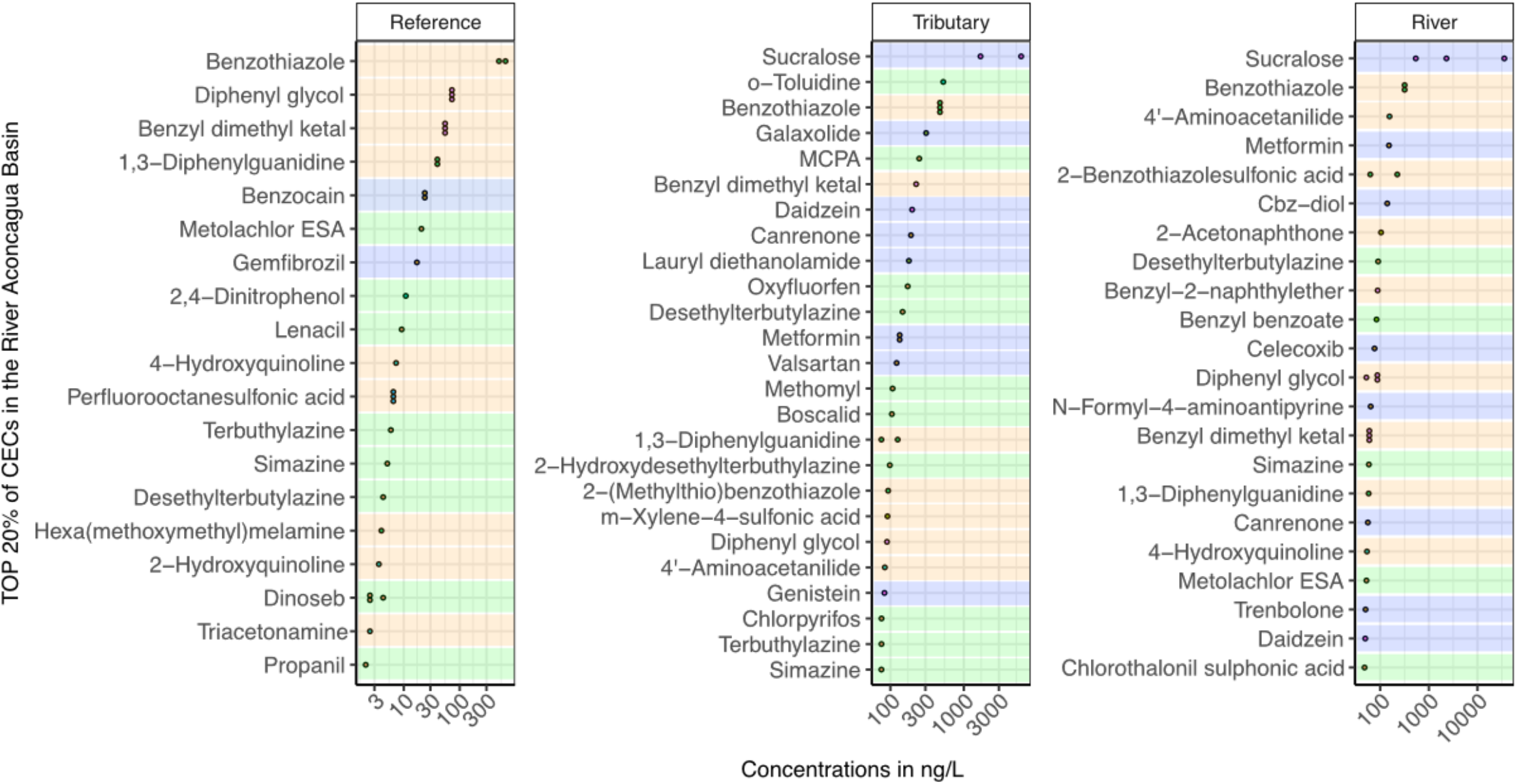
Selected highest concentrations (top 20%) of CECs quantified in at least one sampling site in the River Aconcagua Basin. Main CECs classes are coloured, green represents pesticides, blue PPCPs, and orange industrial chemicals. MCPA = 2-methyl-4-chlorophenoxyacetic acid and Cbz-diol = 10,11-Dihydroxycarbamazepine.

### 3.2. Ecotoxicological assessment

Chronic data for all three main taxonomic groups (algae, macroinvertebrates, and fish) were retrieved for only 34 chemicals (22% of the quantified chemicals) and only for those chemicals the most sensitive taxonomic group can be identified. For a few additional chemicals, we could retrieve partial datasets from the ECOTOX database (Olker et al., 2022), with either data only for algae and macroinvertebrates (7 chemicals), algae and fish (2 chemicals), or macroinvertebrates and fish (2 chemicals). In the end, we are facing the dilemma that chemical-analytical sensitivity and capacity allow screening for hundreds of chemicals of which only a small fraction can be assessed for their risks due to a lack of ecotoxicological data.

In order to evaluate the performance of the QSAR models, we compared the QSAR-estimates to the available experimental chronic data (Figure 2). Unfortunately, all QSAR models show a relatively poor performance (Spearman’s Rho ≤ 0.5) and only ECOSAR predictions are significantly correlated with the experimental data (*p*-value < 0.05) (Figure 2). ECOSAR outperforms VEGA for all three taxonomic groups, showing consistently higher Spearman’s correlation coefficients. Overall, 81% and 85% of the ECOSAR and VEGA predictions, respectively, deviate less than two orders of magnitude from the experimental data. On this basis, we defined nine hazard scenarios for the mixture risk assessment, each with a different strategy to bridge the gaps in the empirical data (Table 1).

**Figure 2.**
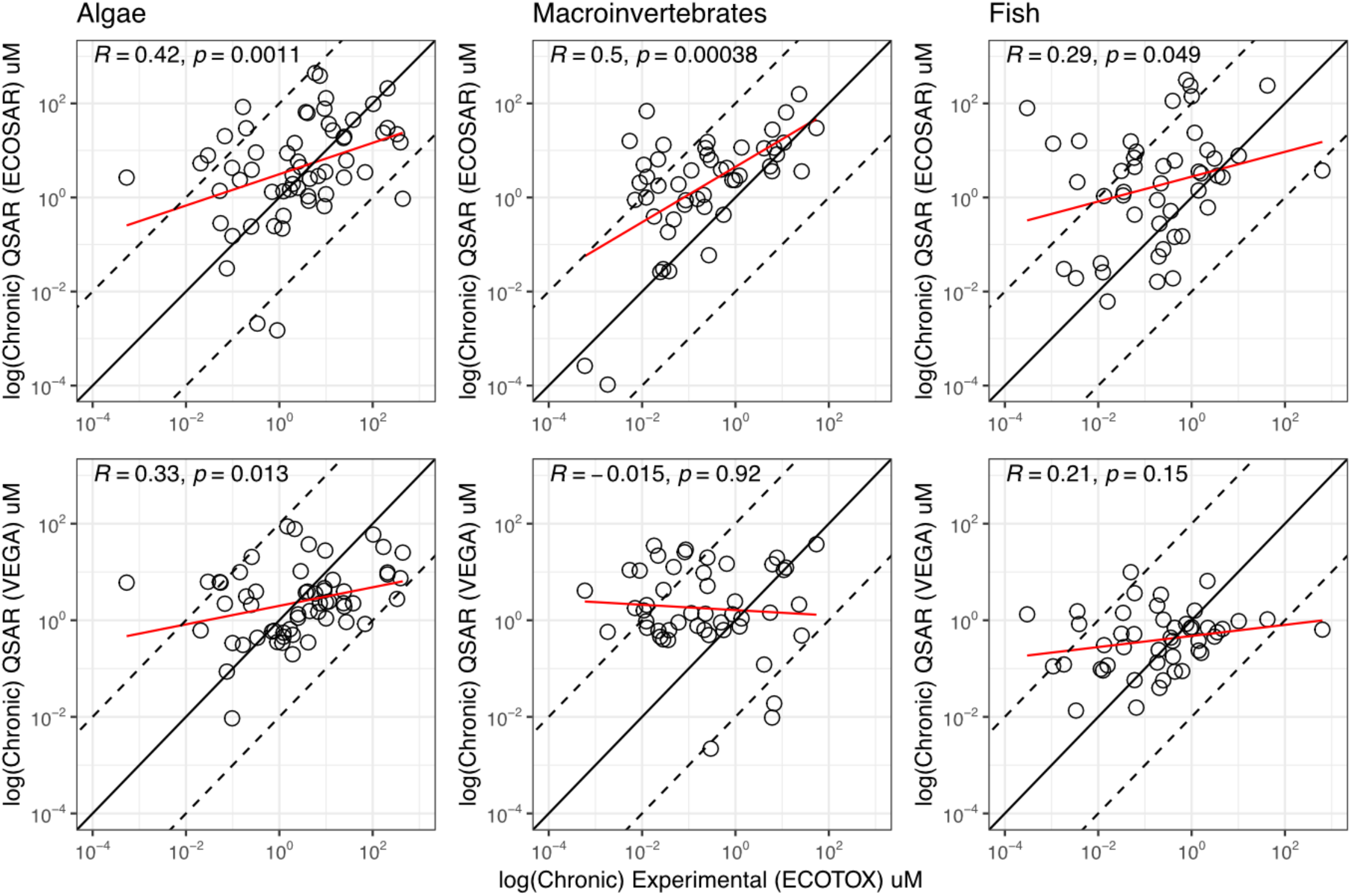
Correlations between experimental and QSAR-based chronic effect data. Upper row shows the results from QSARs estimated using ECOSAR, lower row shows the result from VEGA HUB. Solid lines represent perfect agreement between the experimental and *in silico* predictions and dashed lines denote ± two orders of magnitude deviation. Red lines represent the linear regression model. The non-parametric Spearman’s rank correlation (R) was calculated for significance testing, with the resulting *p*-value provided in each figure.

### 3.3. Mixture risk assessment

In total, we calculated 108 mixture risk scenarios (3 exposure scenarios ξ 9 hazard scenarios ξ 4 mixture evaluations (one for each of the three taxonomic groups plus the MST evaluation)), which were applied to each of the 9 sampling sites included in this study (Figure 3). The resulting 972 mixture evaluations are presented in the supporting information (Table S2). We used a value of 0.1 for the RQ_STU_ and RQ_MST_ sum as the acceptability criterion, which corresponds to applying an assessment factor of 10 to the data, in line with the European Chemical Agency (2016) and the European Commission (2018).

**Figure 3.**
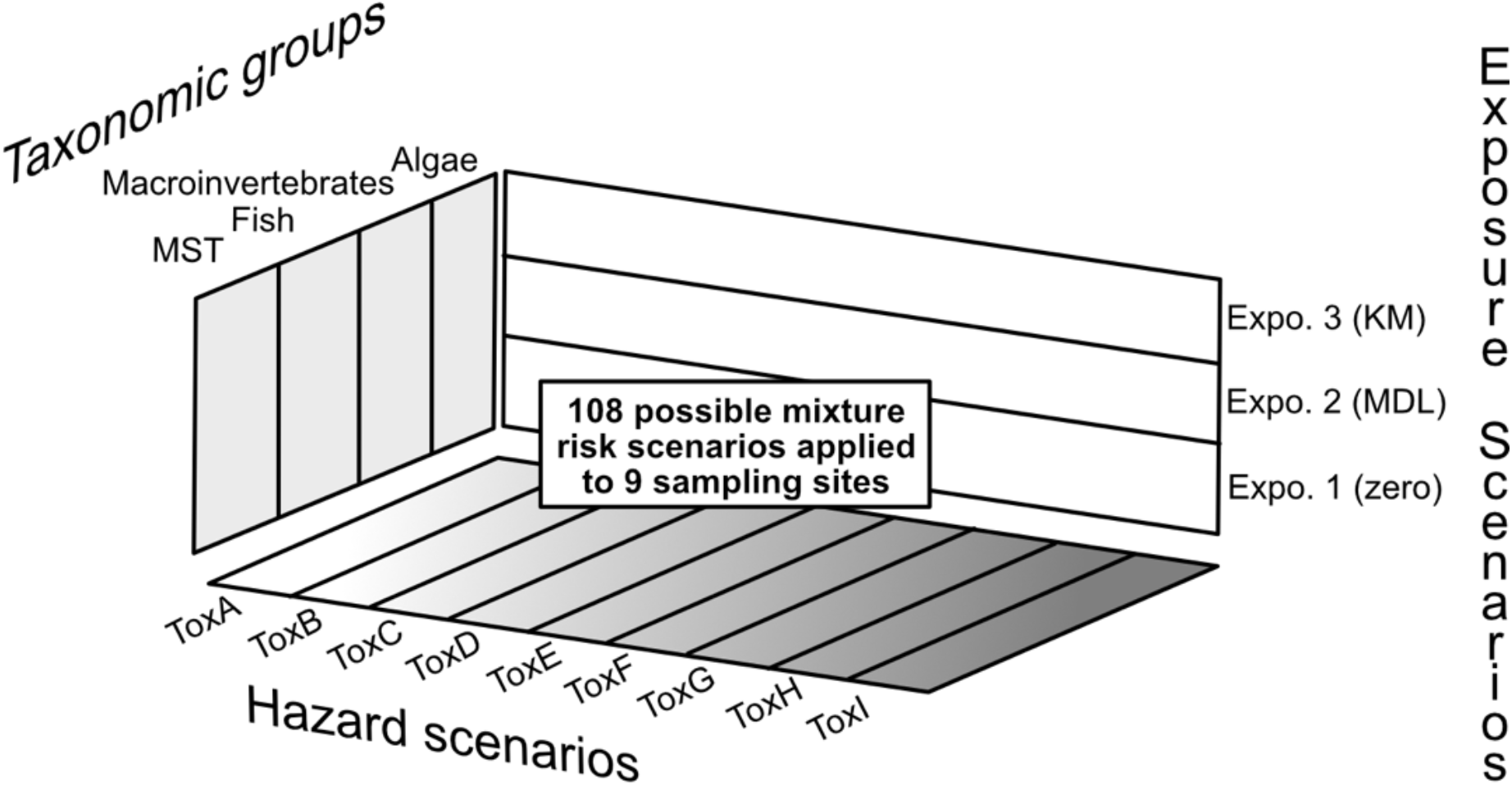
Summary of all mixture risk scenarios applied to the data of each sampling site. Each assessment was performed for each taxonomic group (algae, macroinvertebrates, fish), and the most sensitive taxonomic group (MST). Exposure scenarios are defined based on how non-detects were handled (Expo1 = non-detects set to zero, Expo2 = non-detects set to the MDL, Expo3 = Kaplan-Meier-adjustment), for details see text. The nine hazard scenarios are based on ecotoxicological data used for calculating the risk quotients, see Table 1. A total of 4x3x9 = 108 assessments was calculated for each site.

The Kaplan-Meier-based exposure assessment (Exposure-Scenario 3) always generated risk values that were only marginally higher or lower, respectively, than those generated by Exposure-Scenarios 1 and 2 (ratio of risk estimates between 1.03 and 1.06, Table 3). This shows that the chemical-analytical sensitivity was sufficiently high and that non-detects contribute only marginally to the mixture risk. The Kaplan-Meier scenario was therefore used for the overall mixture risk assessment, while Exposure-Scenario 1 was used for the identification of mixture risk drivers (which cannot be done using Kaplan-Meier estimates, see above). Not surprisingly, risk estimates based on the ToxA scenario (which includes only chemicals with empirical ecotoxicological data) resulted in the lowest risk estimates (Table S2), simply because only a small fraction of the detected chemicals were included. The ToxB and ToxC scenarios (i.e., the mixture assessment based entirely on QSAR-estimates) systematically under-predict mixture risks, in comparison to the corresponding scenarios in which empirical data were preferred (ToxD and ToxG).

**Table 3.** RQ_STU_ estimates for the different exposure scenarios based on the ToxG scenario (empirical chronic data amended with ECOSAR QSAR values). In Expo 1: MECs < MDL were set to 0 (most conservative scenario), Expo 2: MECs < MDL were set to the MDL (worst case scenario), Expo 3: Kaplan-Meier estimation of mixture risk (most realistic scenario). Sites at risk were defined as RQ_STU_ ≥ 0.1, see text. RQ values exceeding 0.1 are boldfaced.

| ToxG (empirical data amended with QSAR-data (ECOSAR)) |  |  |  |  |  |  |  |  |  |  |  |  |
| --- | --- | --- | --- | --- | --- | --- | --- | --- | --- | --- | --- | --- |
|  | Algae |  |  | Macroinvertebrates |  |  | Fish |  |  | MST |  |  |
| Sites | Expo-1 | Expo-2 | Expo-3 | Expo-1 | Expo-2 | Expo-3 | Expo-1 | Expo-2 | Expo-3 | Expo-1 | Expo-2 | Expo-3 |
| RS1 | 1×10<sup>-3</sup> | 0.005 | 0.001 | 0.003 | 0.015 | 0.004 | 0.001 | 0.024 | 0.001 | 0.005 | 0.040 | 0.005 |
| RS2 | 3×10<sup>-4</sup> | 0.004 | 3×10<sup>-4</sup> | 0.001 | 0.013 | 0.001 | 5×10<sup>-4</sup> | 0.024 | 4×10<sup>-4</sup> | 0.001 | 0.037 | 0.001 |
| RS3 | 2×10<sup>-4</sup> | 0.004 | 3×10<sup>-4</sup> | 8×10<sup>-4</sup> | 0.012 | 0.001 | 6×10<sup>-4</sup> | 0.024 | 7×10<sup>-4</sup> | 0.001 | 0.036 | 0.001 |
| T1 | 0.011 | 0.014 | 0.011 | **0.293** | **0.297** | **0.293** | **0.271** | **0.285** | **0.271** | **0.534** | **0.554** | **0.535** |
| T2 | 0.001 | 0.005 | 0.001 | 0.012 | 0.021 | 0.012 | 0.062 | 0.072 | 0.062 | 0.067 | 0.087 | 0.067 |
| T3 | 0.010 | 0.014 | 0.011 | **0.148** | **0.151** | **0.148** | 0.041 | 0.063 | 0.042 | **0.179** | **0.204** | **0.180** |
| R1 | 0.028 | 0.029 | 0.028 | 0.005 | 0.016 | 0.005 | 0.057 | 0.079 | 0.058 | 0.086 | **0.116** | 0.086 |
| R2 | 5×10<sup>-4</sup> | 0.004 | 5×10<sup>-4</sup> | 0.003 | 0.014 | 0.003 | 0.034 | 0.056 | 0.034 | 0.036 | 0.070 | 0.036 |
| R3 | 0.006 | 0.010 | 0.007 | 0.062 | 0.070 | 0.062 | **0.629** | **0.640** | **0.630** | **0.686** | **0.705** | **0.686** |

Because ECOSAR slightly outperformed VEGA for the CECs included in this study (Figure 2), we base the actual mixture risk evaluation on the ToxG scenario (empirical data gaps filled by ECOSAR estimates), see Table 3. The lowest risk was estimated at the reference sites with RQ_Expo3-ToxG-MST_ values ranging between 0.0012 and 0.0056. Tributary streams had higher RQ_Expo3-ToxG-MST_ values, with a site ranking of T1>T3>T2 with RQ_Expo3-ToxG-MST_ values of 0.54, 0.18, and 0.067, respectively. Similarly, the main river course had high RQ_Expo3-ToxG-MST_ values, where sites ranked from R3>R1>R2 with RQ_Expo3-ToxG-MST_ values of 0.69, 0.087, and 0.036, respectively (Table 3). Site R3 (located in the main river course close to the river mouth), site T1 (located in an agricultural area) and site T3 (located in an area with mixed land use) are considered to be at risk (RQ_Expo3-ToxG-MST_ ≥ 0.1).

As expected, RQ_Expo3-ToxG-MST_ values always exceed the corresponding value of the individual taxonomic groups (Table 3). In all three sites where RQ_Expo3-ToxG-MST_ exceeded 0.1 at least one taxonomic group also exceeded a risk quotient of 0.1. That is, the regulatory conclusions from the assessments are identical, independent of whether the mixture risk assessment was performed for each taxonomic group or directly with a view on the whole ecosystem. Values for RQ_Exp3-ToxG-Algae_ were always below the risk threshold of 0.1. We, therefore, consider photosynthetic organisms as not being put at risk by the CEC mixtures analysed in this study. RQ_Exp3-ToxG-Macro_ and RQ_Exp3-ToxG-Fish_ were consistently higher (Table 3).

### 3.4 Mixture risk drivers

We identified eight absolute risk drivers (Table 4). With two exceptions (galaxolide and daidzein), they all belong to groups of substances that are used because of their high biological activity in certain target organisms (pesticides and pharmaceuticals). The top 3 comprise trenbolone, daidzein, and chlorpyrifos with maximum RQ values of 0.62, 0.20, and 0.12, respectively. That is, all three compounds occurred at concentrations that exceed the maximum acceptable level, even if only the exposure to the individual chemical is taken into account (assuming the application of an assessment factor of at least 10, see above).

**Table 4.** Absolute mixture risk drivers, i.e., chemicals with an individual RQ_MST_ ≥ 0.02 in at least one site. Chemicals are listed in descending order of their RQ_Expo1-ToxG-MST_ (mean across all sites). Experimental (ECOTOX) and QSAR ECOSAR concentrations in µM. N = number of sites where the respective chemical was an absolute risk driver.

| Name | CAS Number | Class | minRQ | meanRQ | maxRQ | N | $RQ_{MST-ToxA}$ | $RQ_{MST-ToxC}$ | $RQ_{MST-ToxG}$ | Algae<br>ECOTOX | Macroinv.<br>ECOTOX | Fish<br>ECOTOX | Algae<br>ECOSAR | Macroinv.<br>ECOSAR | Fish<br>ECOSAR |
| --- | --- | --- | --- | --- | --- | --- | --- | --- | --- | --- | --- | --- | --- | --- | --- |
| Trenbolone | 10161-33-8 | Anabolic steroid | 0.045 | 0.330 | 0.615 | 2 | 0.615 | 0.000 | 0.615 |  |  | 0.000 | 16.696 | 5.777 | 79.675 |
| Chlorpyrifos | 2921-88-2 | Insecticide | 0.118 | 0.118 | 0.118 | 1 | 0.118 | 2.057 | 0.118 | 0.248 | 0.001 | 0.012 | 0.240 | 0.000 | 0.025 |
| Daidzein | 486-66-8 | Phytoestrogen | 0.032 | 0.094 | 0.199 | 3 | 0.199 | 0.000 | 0.199 |  |  | 0.003 | 37.523 | 17.450 | 15.898 |
| Galaxolide | 1222-05-5 | Musk fragrance | 0.041 | 0.041 | 0.041 | 1 | 0.041 | 0.062 | 0.041 | 0.777 | 0.028 | 0.398 | 0.245 | 0.030 | 0.019 |
| Diazinon | 333-41-5 | Insecticide | 0.022 | 0.041 | 0.061 | 2 | 0.061 | 0.139 | 0.061 | 9.189 | 0.000 | 0.433 | 0.653 | 0.000 | 0.144 |
| Methomyl | 16752-77-5 | Insecticide | 0.036 | 0.036 | 0.036 | 1 | 0.036 | 0.001 | 0.036 | 440.749 | 0.018 | 0.183 | 0.940 | 0.393 | 0.882 |
| Terbuthylazine | 5915-41-3 | Herbicide | 0.021 | 0.029 | 0.037 | 2 | 0.021 | 0.000 | 0.021 | 0.052 | 0.008 | 3.569 | 1.384 | 2.062 | 2.909 |
| Clarithromycin | 81103-11-9 | Antibiotic | 0.026 | 0.026 | 0.026 | 1 | 0.026 | 0.000 | 0.026 | 0.000 |  |  | 2.650 | 1.453 | 1.266 |

We also determined 19 substances as *potential* absolute risk drivers (Table S3). Those are substances without a full set of empirical ecotoxicity data, but which could potentially be risk drivers, under the worst-case assumption that their QSAR-based hazard estimate underestimates their actual toxicity by a factor of 100 (see Figure 2 and Table 4 for the ratios between empirical data and QSAR estimates). Twelve substances did not have any empirical data on chronic toxicity, five chemicals had chronic information for one taxonomic group, and only two chemicals (i.e., fungicides boscalid and myclobutanil) have chronic information for two taxonomic groups. Especially relevant are pesticides, biocides and pharmaceuticals with a maximum RQ_Expo1-ToxI-MST_ ≥ 0.1 for which either the experimental data on the likely target organism (= the most sensitive organism group) are missing, such as chlorfenapyr (an insecticide), telmisartan (a pharmaceutical) and allethrin (an insecticide); or chemicals for which we don’t know their target organisms, such as octocrylene (a personal care product), benzyl-2-naphthyl ether (an industrial chemical), or N,N-Dimethyltetradecylamine N-oxide (TDAO, a surfactant). Empirical ecotoxicological data are urgently needed for these substances.

Eight relative risk drivers were determined across the nine sites (Table S4). All of them, with two exceptions, were also categorised as actual absolute risk drivers. The two exceptions (1,3-diphenylguanidine and chlorfenapyr) were also identified as *potential* absolute risk drivers. We also identified six chemicals as *potential* relative risk drivers (Table S5) of which four were also categorised as *potential* absolute risk drivers.

The mixture risk drivers at those sites where risk cannot be excluded (RQ_Expo1-ToxI-MST_ ≥ 0.1) are shown in Figure 4 and in those with RQ_Expo1-ToxI-MST_ < 0.1 (no risk) are presented in Figure S2. All three sites at risk (i.e., T1, T3, and R3) show distinct patterns and possess different risk drivers (Figure 4). The number of absolute as well as relative risk drivers never exceeded 4. This is a typical pattern in environmentally realistic mixtures, which is sometimes called the Pareto-principle of mixture toxicity, relating to the power-law probability distribution named after the Italian engineer Vilfredo Pareto (Rudén et al., 2019 and references therein). However, the chemicals actually identified as risk drivers varied across sites, in dependence on land-use patterns and land-use intensity.

**Figure 4.**
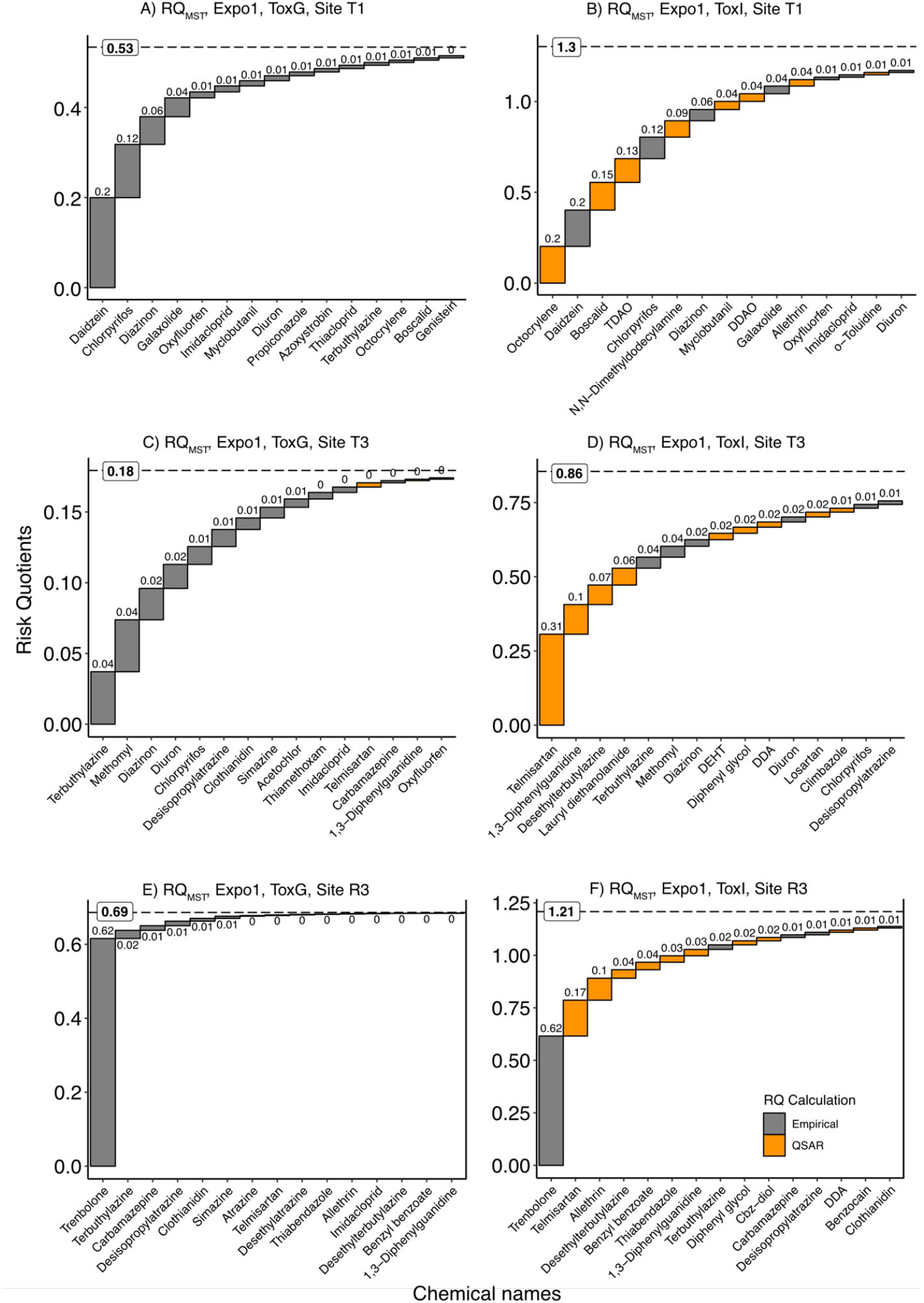
Mixture risk assessment estimates for the sites at risk (RQ_MST_≥0.1) in the River Aconcagua basin. (A)(C)(E): RQ_MST_ predictions based on non-detects set to zero (Expo1) and experimental chronic data amended with ECOSAR values (ToxG). (B)(D)(F): RQ_MST_ predictions based on non-detects set to zero (Expo1) and experimental chronic data amended with ECOSAR values (ToxG) that were multiplied by a factor of 100 (i.e., QSAR ECx estimates were divided by a factor of 100) (ToxI). Colours represent the source of the toxicity data. Grey bars: empirical data, orange data: QSAR estimates.

## 4. Discussion

### 4.1. Exposure and hazard assessment

The nature of the chemicals found and the fact that we observed a clear pollution gradient from the “reference sites” to the main river and the tributaries shows the impact of human activities on the chemical status of the River Aconcagua basin. The overall CEC fingerprints did not substantially differ from those previously determined in Europe, North America, and some African countries (Carpenter and Helbling, 2018; Finckh et al., 2022; Kandie et al., 2020; Loos et al., 2013). This similarity can be attributed to the widespread and global use of these chemicals in daily life, industry, and agriculture, as well as the used of a target list based on commonly measured CECs in European aquatic environments. The CECs that we detected at the highest concentrations, sucralose and benzothiazole, are ubiquitous in surface waters around the globe, in similar concentration ranges (Finckh et al., 2022; Loos et al., 2013; Yu et al., 2023).

Although we have the analytical sensitivity for screening hundreds of CECs in the aquatic environment, the Achilles’ heel is the lack of ecotoxicological data for assessing CEC hazards. A similar situation has been described in previous publications, including studies that assessed WWTP effluents (Finckh et al., 2022), agricultural streams after rain events (Neale et al., 2020), and emission-based mixture risk assessments (Gustavsson et al., 2023). The use of chronic effect estimates derived from QSARs bridges the gap in chronic effect data, which enabled us to conduct the mixture assessment separately for each of the three taxonomic groups (algae, macroinvertebrates, fish) and MST for all the CECs included in our study. Nevertheless, our results show that the accuracy of the chronic QSAR estimates needs improvement, findings that are in agreement with previously published studies, with some exceptions in the field of endocrine disruption (Cronin, 2017). QSAR models that estimate acute ecotoxicity perform better (Melnikov et al., 2016; Zhou et al., 2021).

### 4.2. Mixture risk assessment and risk drivers

Our study presents a systematic and comprehensive strategy for the environmental risk assessment of chemical mixtures. This strategy encompasses 108 distinct mixture risk scenarios, taking into account different possibilities on how to account for the potential contribution of CECs that were not detected, different strategies to account for data gaps and different ecotoxicological perspectives (focus on individual taxonomic groups and MST). The role of non-detects has only rarely been systematically evaluated within the framework of mixture assessment, and only a limited number of studies have accounted for their potential contribution to the mixture risk (Gustavsson et al., 2017a, 2017b). Conversely, the incorporation of QSAR predictions into the mixture assessment framework is generally applied (Finckh et al., 2022; Reiber et al., 2021; von der Ohe et al., 2011), as recommended by the European Chemical Agency (European Chemicals Agency, 2021). In addition to using QSARs for filling data gaps, we introduce a novel application of QSARs to predict *potential* mixture risk drivers. This approach identifies chemicals that may have adverse effects at the concentrations at which they are detected and for which therefore empirical data are urgently needed.

Three of the nine sites can be considered at risk (Sum RQ ≥ 0.1 in scenario ToxG, using KM-based exposure estimates) in the River Aconcagua basin. The small number of risk drivers found at each of those sites raises the question of whether the risk is indeed an issue that is driven by mixtures, or whether this is a single-substance problem, and the analytical performance was simply good enough to detect a myriad of chemicals that happen to co-occur but that are irrelevant from a risk perspective. If risk mitigation measures would focus exclusively on those compounds that are present at unacceptable concentrations (individual RQ > 0.1) in order to reduce their individual RQ values to a maximum of 0.1, the remaining RQ sums would be 0.23 (site T1), 0.18 (site T3) and 0.17 (site R3). That is, even if single-substance-oriented risk mitigation measures would be consistently implemented so that all chemicals are present at individually “safe” concentrations, all three sites would still be at risk. This leads to the conclusion that the risks encountered at the sites are a combination of a single substance problem (unacceptably high concentrations of a few individual substances), and a mixture problem (unacceptably high sum RQ values even after successful single substance risk mitigation).

Component-based mixture risk assessments, such as the one implemented in this study, inherently underestimate the actual site-specific risks, given that most likely not all relevant chemicals are included in the analytical profile. For instance, the present study is based on a selection of target compounds that was developed largely from a European perspective (Beckers et al., 2018; Krauss et al., 2019), and therefore does not include all pesticides used in Chilean agriculture. The present study also focuses exclusively on environmental pollution by synthetic organic chemicals and overlooks the role of metals as risk-contributing contaminants. At the same time, the assumption of a concentration-additive behaviour of the mixture might lead to a risk overestimation, although this might be comparatively small (see discussion in Backhaus and Faust, 2012; Rudén et al., 2019).

The top absolute risk drivers (RQ_MST_ ≥ 0.02, Table 4) are CECs that are known to cause harmful effects on aquatic life. Trenbolone stands out as a veterinary drug that enters river basins through livestock farming, which has been recognised as an endocrine disruptor, capable of altering hormone and steroid synthesis in fish (Ankley et al., 2008; Overturf et al., 2015). Chlorpyrifos is a chlorinated organophosphate insecticide that is well-known for its neurotoxicity to invertebrates and fish (Echeverri-Jaramillo et al., 2020; Scott and Sloman, 2004). Daidzein is a natural phytoestrogen primarily found in the *Fabaceae* family, including soybeans, peas, and red clover. Chlorpyrifos and daidzein have been previously identified as risk drivers in the aquatic environment (Caracciolo et al., 2023; König et al., 2017). In addition, the absolute risk drivers diazinon, terbuthylazine, and clarithromycin have been also identified as major risk drivers in WWTP’s effluents (Beckers et al., 2018; Finckh et al., 2022). Chlorpyrifos is a priority substance of the EU Water Framework Directive and the sunscreen octocrylene, identified as a *potential* absolute risk driver, is listed on the 3^rd^ watch list under the WFD (European Commission, 2022, 2008).

## 5. Conclusions

Our study advances our understanding of environmental risks caused by the co-occurrence of CECs in freshwater systems in South America. In the River Aconcagua basin, we detected a total of 153 CECs from various chemical classes, including pesticides, pharmaceuticals, personal care products, and chemicals used in industrial processes. The overall pattern of CEC occurrence did not differ significantly from other small streams and rivers worldwide. However, we observed clear site-specific differences in concentrations and mixture composition.

To comprehensively evaluate the risk associated with CEC mixtures, we introduced an integrative strategy for mixture risk assessment. This approach systematically assesses both the exposure and hazard components of the risk assessment process. Due to the lack of experimental ecotoxicological data, we utilized QSAR modelling, as recommended by various environmental agencies, to fill data gaps. The QSAR models lacked accuracy across different taxonomic groups, and we incorporated those uncertainties into the mixture risk assessment, by defining various hazard scenarios.

Based on our analysis, we identified three sites at risk in the River Aconcagua basin. These conclusions are supported by the different risk scenarios and their interlinkage. Furthermore, our findings endorse the use of the most sensitive taxonomic group (RQ_MST_) as a comprehensive ecological risk metric for predicting the risks posed by complex environmental mixtures. This metric successfully captured the taxonomic groups that were most vulnerable to the determined exposures.

Risk scenarios based solely on QSAR ecotoxicological data consistently underestimated the actual risk. Therefore, we propose the use of QSAR predictions amended with experimental ecotoxicological data as a worst-case scenario for risk estimation. QSAR models proved to be valuable for identifying chemicals that potentially contribute to the predicted risk (*potential* risk drivers). Additionally, we recommend evaluating the performance of available QSAR platforms, especially those offering chronic models, before integrating their predictions into the risk assessment process.

We found that only a few chemicals were responsible for driving the mixture risk. However, the results show that mitigation measures focused solely on single chemicals are insufficient if water bodies are impacted by complex mixtures of chemicals. It is crucial to acknowledge that chemical pollution risks are a combination of (1) the problem of unacceptably high concentrations of comparatively few individual substances and (2) the problem caused by a complex melange of chemicals, co-occurring at seemingly low concentrations.

## 6. CRediT authorship contribution statement

**Pedro A. Inostroza**: Conceptualization, Methodology, Software, Validation, Formal analysis, Investigation, Data curation, Writing – Original Draft, Writing – review & editing, Visualisation. **Sebastian Elgueta:** Methodology, Writing – review & editing. **Martin Krauss**: Methodology, Data curation, Writing – review & editing. **Werner Brack**: Resources, Writing – review & editing. **Thomas Backhaus**: Conceptualization, Methodology, Software, Validation, Formal analysis, Investigation, Data curation, Writing – Original Draft, Writing – review & editing, Visualisation, Funding acquisition.

## 7. Acknowledgments

We thank Alba Lopez Mangas and Monica del Aguila for fieldwork support, Margit Petre and Jörg Ahleim (UFZ) for LVSPE training as well as cartridge extractions. This work was supported by the FRAM Centre for Future Chemical Risk Assessment and Management at the University of Gothenburg. The QExactive Plus LC-HRMS used at UFZ is part of the major infrastructure initiative CITEPro (Chemicals in the Terrestrial Environment Profiler) funded by the Helmholtz Association.

